# Impact of RNA Extraction and Target Capture Methods on RNA Sequencing Using Formalin-Fixed, Paraffin Embedded Tissues

**DOI:** 10.1101/656736

**Authors:** Christopher A. Hilker, Aditya V. Bhagwate, Jin Sung Jang, Jeffrey G Meyer, Asha A. Nair, Jaime I. Davila, Amber M. McDonald, Jennifer L. Winters, Rebecca N. Wehrs, Rory A. Jackson, Joshua A. Gorman, Mine S. Cicek, Andre M. Oliveira, E. Aubrey Thompson, Bruce W. Eckloff, Kevin C. Halling, Zhifu A. Sun, Jin Jen

## Abstract

Formalin fixed paraffin embedded (FFPE) tissues are commonly used biospecimen for clinical diagnosis. However, RNA degradation is extensive when isolated from FFPE blocks making it challenging for whole transcriptome profiling (RNA-seq). Here, we examined RNA isolation methods, quality metrics, and the performance of RNA-seq using different approaches with RNA isolated from FFPE and fresh frozen (FF) tissues. We evaluated FFPE RNA extraction methods using six different tissues and five different methods. The reproducibility and quality of the prepared libraries from these RNAs were assessed by RNA-seq. We next examined the performance and reproducibility of RNA-seq for gene expression profiling with FFPE and FF samples using targeted (Kinome capture) and whole transcriptome capture based sequencing. Finally, we assessed Agilent SureSelect All-Exon V6+UTR capture and the Illumina TruSeq RNA Access protocols for their ability to detect known gene fusions in FFPE RNA samples. Although the overall yield of RNA varied among extraction methods, gene expression profiles generated by RNA-seq were highly correlated (>90%) when the input RNA was of sufficient quality (≥DV_200_ 30%) and quantity (≥ 100 ng). Using gene capture, we observed a linear relationship between gene expression levels for shared genes that were captured using either All-Exon or Kinome kits. Gene expression correlations between the two capture-based approaches were similar using RNA from FFPE and FF samples. However, TruSeq RNA Access protocol provided significantly higher exon and junction reads when compared to the SureSelect All-Exon capture kit and was more sensitive for fusion gene detection. Our study established pre and post library construction QC parameters that are essential to reproducible RNA-seq profiling using FFPE samples. We show that gene capture based NGS sequencing is an efficient and highly reproducible strategy for gene expression measurements as well as fusion gene detection.

---

RNA Sequencing (RNA-seq) is increasingly used to provide a complete profiling of the entire transcriptome in a given sample at the resolution of individual transcripts. Like most RNA analysis methodologies, RNA-seq performs best when using high quality RNA isolated from fresh frozen tissue samples. Since less than 5% of total cellular RNA is comprised of protein coding mRNA (1), genomic analysis by RNA-seq generally focuses on gene coding mRNA isolated via poly-A sequences common to the 3’ end of the transcript. When RNA is degraded, this poly-A based selection approach results in a biased preference for the sequencing of the 3’ ends of the transcript (2-4).

In clinical situations, FFPE samples are the most commonly used but they are challenging for molecular analysis due to extensive nucleic acid degradation that results from formalin fixation (5). To overcome this limitation, an alternative approach for RNA-seq with FFPE specimens is to enrich mRNA by reducing the ribosomal RNA (rRNA) fraction in the sample using oligo nucleotides with complementary rRNA sequences or enzymes that specifically degrade rRNA (6, 7). More recently, direct hybridization of mRNA through targeting and capturing only genes of interest has shown great promise, particularly for partially degraded RNA from FFPE samples (8, 9). Several commercial kits are now available to capture a preselected set of RNA targets or the entire transcriptome from FFPE RNA samples (Supplementary Data Sheet Publications).

In this study, we evaluated commonly used FFPE RNA isolation and targeted RNA capture methods with the aim to develop a reliable protocol for whole transcriptome profiling by Next Generation Sequencing (NGS) using archival tissues.

## Materials and Methods

### Samples and Methods Used for RNA Extraction Evaluation

We used six different archival tissue blocks (liver, colon (n=2), pancreas, kidney, and tonsil) to evaluate the average performance of five different RNA extraction methods (Table1). Under the protocol approved by the Institutional Review Board, one millimeter cores (one core per sample for each extraction) were taken from each of the six FFPE blocks and RNA was isolated using High Pure FFPE RNA Micro (Roche, Basel, Switzerland), miRNeasy FFPE and RNeasy FFPE (Qiagen, Hilden, Germany) and RecoverAll Total Nucleic Acid Isolation (Ambion, Austin, TX) kits following manufacturers’ protocols. Total RNA recoveries were quantified with a NanoDrop instrument (Thermo Fisher Scientific, Waltham, MA) and fragment sizes were analyzed using a 2100 Bioanalyzer (Agilent Technologies, Santa Clara, CA). TruSeq Stranded Total RNA Library Prep (Illumina, San Diego, CA) method was used to construct the sequencing libraries following the manufacture’s protocol (Supplementary Methods).

### Samples Used to Assess RNA-Seq by Target Capture

We next used a genetically engineered cell line T47D+ which contains an *ESR1-YAP1* fusion introduced into its parental cell line (T47D-) to evaluate overall gene expression correlation and fusion gene detection between whole transcriptome capture by SureSelect All-Exon (v4 + UTR) and TruSeq mRNA. Matched FFPE and FF samples from a phosphaturic mesenchymal tumor (PMT) and a synovial sarcoma (SC1) were used to evaluate gene expression correlation between targeted and whole transcriptome capture for FFPE RNA, and between whole transcriptome capture of the FFPE RNA and poly-A selection of the matched FF RNA by mRNA-seq (Table 1). Clinically significant gene fusions were then evaluated using FF RNA samples (R106 normal kidney, R130 Diffuse Large B-Cell Lymphoma and R152 colorectal cancer), the matched FFPE samples (R106 FFPE, R130 FFPE) and an additional FFPE colorectal cancer sample (R153). All RNA samples were extracted using either Qiagen miRNeasy or the Qiagen miRNeasy FFPE kits following manufacture’s protocols, for FF and FFPE samples, respectively.

**Table 1.** Summary of Experimental Studies and Samples.

| Purpose | Samples | Library Preparation Method | Capture Method |
| --- | --- | --- | --- |
| **RNA extraction comparison** | • Normal Tonsil (FFPE)<br>• Liver Tumor (FFPE)<br>• Colon Tumors (2) (FFPE)<br>• Pancreas Tumor (FFPE)<br>• Kidney Tumor (FFPE) | • TruSeq Stranded Total RNA (Illumina) | • RiboZero* |
| **Gene Capture for Fusion Detection** | • T47D +/- Fusion | • TruSeq Stranded mRNA (Illumina) | • SureSelect All-Exon v4 + UTR<br>• PolyA |
| **All-Exon vs Kinome Capture vs TruSeq mRNA** | • PMT (FF/FFPE)<br>• SC1 (FF/FFPE) | • NEBNext Ultra RNA w/ capture bait.<br>• TruSeq mRNA (Illumina) | • SureSelect All-Exon v4 + UTR (FFPE)<br>• SureSelect Kinome (FFPE)<br>• PolyA (FF) |
| **Illumina Access vs Agilent SureSelect Capture** | • R106 (FF/FFPE)<br>• R130 (FF/FFPE)<br>• R152 (FF)<br>• R153 (FFPE) | • TruSeq Access (Illumina)<br>• SureSelect RNA for Target Enrichment (Agilent) | • TruSeq Access<br>• SureSelect All-Exon v6 +UTR |

\*RiboZero captures and eliminates rRNA prior to library preparation

### RNA Library Preparations and Capture

Library preparation methods used for each comparison study are summarized in Table 1 and were carried out following the manufactures’ protocols (Supplemental Methods). Briefly, 100-150 ng of total RNA samples were used for cDNA synthesis after rRNA depletion (Stranded Total RNA), poly-A RNA selection (Stranded and TruSeq mRNA) or without selection (NEBNext Ultra, TruSeq Access and SureSelect RNA Target Enrichment kits). For gene capture involving SureSelect baits, pre-capture adapters (Agilent) were included in the library preparation step. Pre-capture libraries were quantified by Qubit (Thermo Fisher Scientific, Waltham, MA) and a 2100 Bioanalyzer, target-captured using Kinome, All-Exon, or TruSeq Access baits (S Table 1), purified with AMPure XP beads (Beckman Coulter, Brea, CA) and then PCR amplified following manufacturer’s recommendation. Detailed methods for each library preparation are available upon request. Post-capture libraries were quantified again by Qubit and a 2100 Bioanalyzer prior to sequencing on a HiSeq 2000/2500 (101 cycles and paired end reads at 5-8 samples per lane).

### Bioinformatics Data Analysis and Assessment

The raw RNA-seq data in fastq format was processed by the Mayo Bioinformatics Core pipeline using our MAP-RSeq v.1.3.0.1 workflow (10). MAP-RSeq consists of alignment with TopHat 2.0.6 (11) against the human hg19 genome build and gene expression quantification with the Subread package 1.4.4 [PMID: 24227677] against RefSeq annotation, which was obtained from Illumina (http://cufflinks.cbcb.umd.edu/igenomes.html). Gene expression of the raw read counts was normalized using the RPKM (Reads per Kilobase per million Mapped reads) approach. The log2 transformed gene expression data was used to conduct pair-wise correlation by Pearson correlation coefficient between RNA extraction protocols, capture methods, and FF and FFPE samples. MAPR-Seq also conducts expressed fusion transcript detection using the TopHat-Fusion algorithm (12). Gene fusions detected by Tophat-Fusion are then processed through an in-house fusion annotation module that specifically scans for fusions occurring in a focused set of 573 cancer-related genes (12). To consider a sample positive for a gene fusion, the module requires a minimum of 5 fusion supporting reads, greater than 100kb distance between the fusion breakpoints and that the fusion has to be at exon-exon boundaries of the fusion gene pairs. These fusions were also visually examined by loading the alignment files for contrast protocols into the Integrative Genomics Browser (IGV) (13). Differences in probe designs between the different capture methods were also inspected in IGV.

## Results

### Impact of RNA Yield and Quality of RNA Extraction Methods on RNA-seq

We used six different human tissues to provide sample diversity in assessing the overall performance of commonly used RNA isolation methods for FFPE samples. We also used average values when comparing RNA yields and QC metrics. Among the samples tested using four different RNA isolation kits, the RecoverAll FFPE kit gave the highest average RNA yield at 10.1 µg for the six samples. The average total RNA yield with the other kits was lower with RNeasy FFPE at 7.4 µg, miRNeasy FFPE at 8.9 µg, miRNeasy on Qiacube at 5.9 µg and High Pure at 2.5 µg (Table 2). The RNA yield for each individual sample is shown in S Table 2.

**Table 2.** FFPE RNA extraction kit comparison.

| Kit Name | Total Yield/Sample*<br>(µg) (SD) | 260/280<br>(SD) | 260/230<br>(SD) | RIN<br>(SD) | DV200**<br>(SD) |
| --- | --- | --- | --- | --- | --- |
| RecoverAll Total Nucleic Acid Isolation Kit (Ambion) | 10.1<br>(4.1) | 1.98<br>(0.06) | 1.61<br>(0.40) | 2.4<br>(0.08) | 61.67%<br>(12.83%) |
| RNeasy FFPE Kit (Qiagen) | 7.4<br>(2.4) | 1.95<br>(0.04) | 1.96<br>(0.15) | 2.5<br>(0.12) | 37.83%<br>(15.77%) |
| miRNeasy FFPE Kit (Qiagen) | 8.9<br>(5.7) | 1.93<br>(0.03) | 1.93<br>(0.12) | 2.5<br>(0.11) | 30.83%<br>(20.52%) |
| miRNeasy FFPE Kit on Qiacube (Qiagen) | 5.9<br>(2.9) | 1.94<br>(0.04) | 1.86<br>(0.14) | 2.5<br>(0.10) | 50.33%<br>(8.69%) |
| High Pure FFPE RNA Micro Kit (Roche) | 2.5<br>(1.6) | 1.79<br>(0.23) | 1.05<br>(0.69) | 2.3<br>(0.20) | 73.00%<br>(4.38%) |

\* The average values based on all 6 samples used for each extraction method, with standard deviation (SD), and calculated from either NanoDrop (total yield, 260/280, and 260/230), or Bioanalyzer (RIN and DV200).
\*\* DV<sub>200</sub> values reflect the percentage of RNA fragments greater than 200nt in size.

Using absorbance ratios at 260/280 and 260/230nm values as purity measures, the three Qiagen methods averaged between 1.93-1.95 and 1.86-1.96; the RecoverAll had an average of 1.98 and 1.61, while samples isolated using the High Pure Kit had an average of 1.79 and 1.05, respectively (Table 2). Although the same tissues were used, the average fraction of fragment sizes above 200 bases (DV_200_ values) for the isolated RNA samples varied significantly among different extraction methods, in part reflecting sample purity as well as efficiency of RNA recovery. For example, although the High Pure kit resulted in RNA samples with the highest average DV_200_ value of 73%, the lower purity values suggest that non-RNA contaminants likely contributed to the DV_200_ readings. In contrast, Qiagen kits tended to have lower DV_200_ values as well as lower total RNA yield. Not unexpectedly, RNA Integrity (RIN) values were low (≤2.5) for all tested FFPE RNA samples.

Using the Illumina TruSeq stranded RNA-seq method, the total reads, reads mapped to the genome, mapped exon-exon junction and gene counts were comparable across samples and three extraction methods (S. Table 3). Significantly, gene expression correlations for the same samples extracted by different methods showed high correlations for a majority of the tested samples (examples in Figure 1, R = 0.92 - 0.97).

**Table 3.**
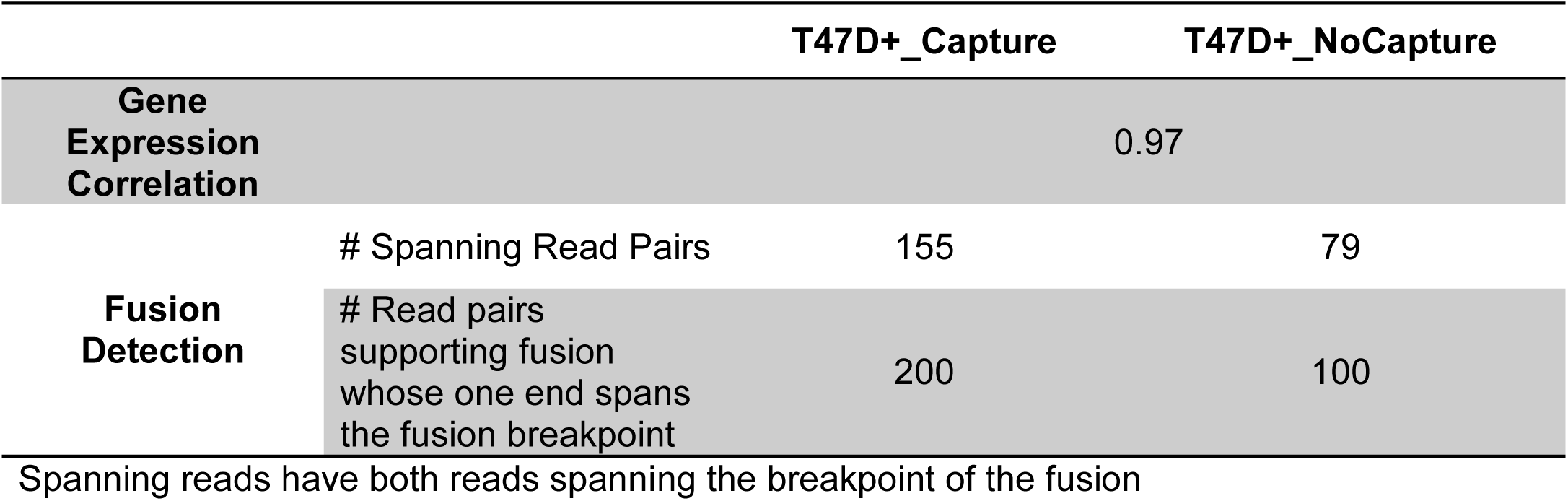
Fusion Supporting Reads for Sample T47D+ with and without All-Exon Capture.

**Fig 1.**
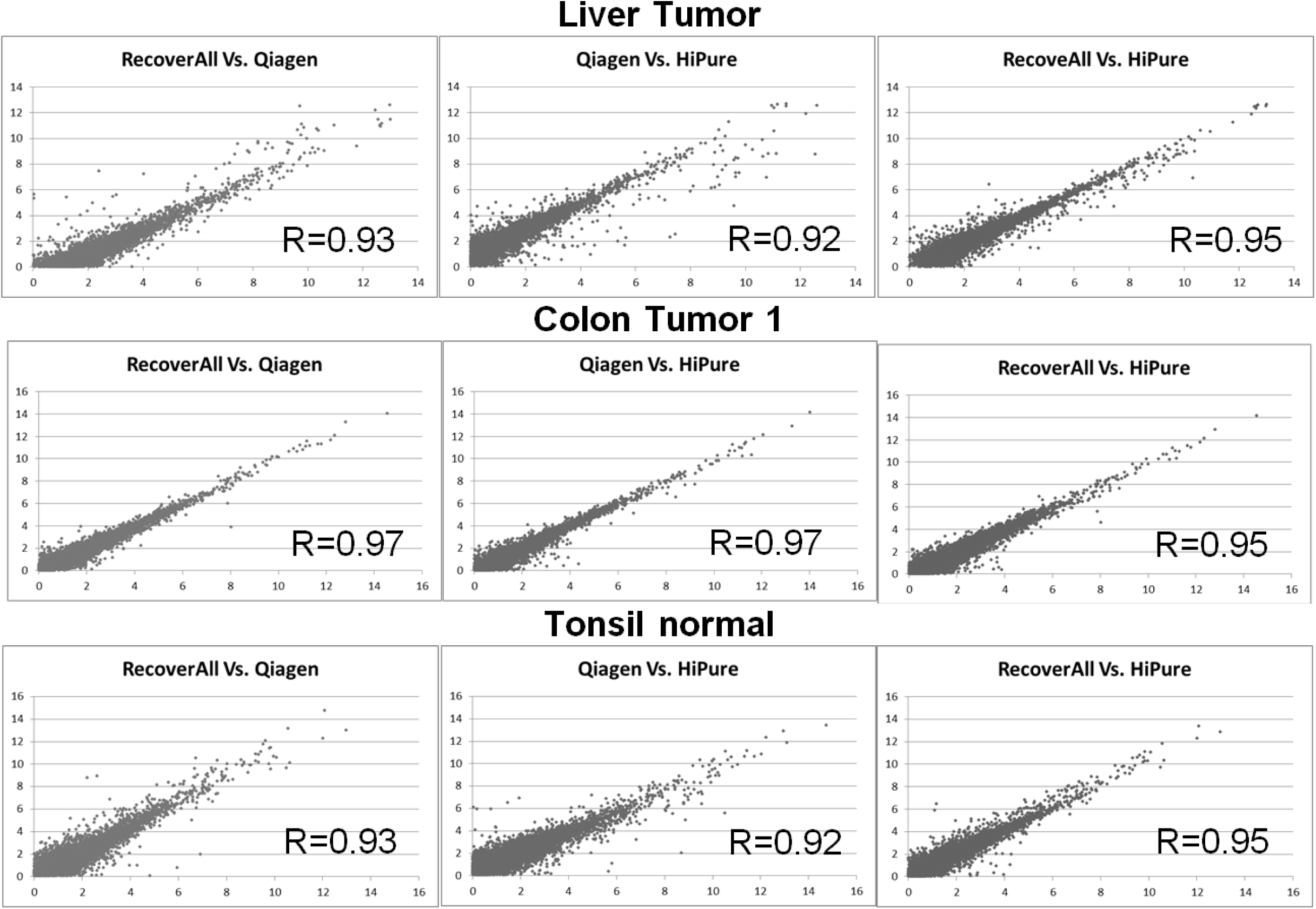
Impact of RNA Extraction Method on RNA-seq Gene Expression Measurements. Representative gene expression correlations among FFPE RNA samples that were extracted us**i**ng RecoverAll, HiPure and Qiagen miRNeasy FFPE extraction kit

### Fusion Transcript Detection by Gene-specific Capture vs. poly-A Selection

We next assessed whether gene-specific capture using Agilent SureSelect baits (All-Exon v4 + UTR) could enhance fusion gene capture as compared to poly-A selection by using two cell lines with and without an engineered *ESR1-YAP1* fusion gene (T47D+ and T47D-). Overall, gene coverage between the two samples and methods was very similar with 17,386-17,958 genes at ≥1X coverage and 15, 981-16,361 genes when ≥ 5X coverage were considered (S Table 4). However, the number of reads supporting the fusion was two times greater with the gene capture protocol than traditional poly-A selection and TruSeq mRNA prep. Gene expression correlation for these two samples was high between gene-specific capture and poly-A selection methods(R = 0.97) (Table 3). Observed reads for the fusion breakpoint in sample T47D+ had significantly higher spanning read pairs by gene-specific capture (155) as compared to those obtained by poly-A selection (79) (Table 3). Read pairs with one end spanning the fusion breakpoint also were higher by the gene capture method (200 reads) compared to 100 reads with poly-A selection (Table 3).

**Table 4.** Fusion Detection in Fresh Frozen and FFPE Tissue using TruSeq mRNA, Access Capture and SureSelect Capture.

**A - Fresh Frozen Tissue**
| Sample | RIN | Dist. To Poly A | Fusion | Reads Supporting Fusion |  |  |
| --- | --- | --- | --- | --- | --- | --- |
|  |  |  |  | TruSeq mRNA | Access Capture | SureSelect Capture |
| R106 | 7.2 | -- | None | 0 | 0 | 0 |
| R130 | 9.5 | >5kb | *LPP-FOXP1* | 0 | 11 | 7 |
| R130 | 9.5 | >15kb | *FOXP1-LPP* | 0 | 40 | 4 |
| R152 | 7.6 | >5kb | *FGFR2-BICC1* | 7 | 605 | 121 |
| R152 | 7.6 | 1.3kb | *BICC1-FGFR2* | 113 | 111 | 27 |

**B - FFPE Tissue**
| Sample | RIN | Dist. To Poly A | Fusion | Reads Supporting Fusion |  |
| --- | --- | --- | --- | --- | --- |
|  |  |  |  | Access Capture<br>FFPE miRNeasy | SureSelect Capture<br>FFPE miRNeasy |
| R106 | 2.4 | -- | None | 0 | 0 |
| R130 | 2.4 | >5kb | *LPP-FOXP1* | 5 | 0 |
| R130 | 2.4 | >15kb | *FOXP1-LPP* | 14 | 0 |
| R153 | 2.4 | 3.8kb | *FGFR2-WAC* | 368 | 57 |
| R153 | 2.4 | 3.2kb | *TCF7L2-FAM184B* | 30 | 18 |

Fusion supporting reads for FF and FFPE samples with known fusion.

### Gene Expression Correlation by Targeted Gene Capture vs. All-Exon Capture

We investigated whether the size and complexity of the target capture region affect the performance of gene expression measurements using matched FF and FFPE tissues (PMT and SC1). Two SureSelect bait sets, Kinome (3.16 Mb) and All-Exon V4+UTR (71.45 Mb), were used for gene expression correlation evaluation (S Table 1). At a sequencing depth of approximately 36 million reads per sample, 92.9%-93.8% of the reads were mapped to specific reference sequences for FFPE RNA while a slightly higher fraction mapped (95.9%-96.2%) to specific reference sequences using RNA from FF tissues (S Table 5). As expected, the average base coverage for the Kinome captured samples was significantly higher than for All-Exon captured samples for FFPE samples, but did not differ greatly in FF samples (S Figure 1). To directly compare the two capture bait sets for gene expression quantification, we focused on gene expression correlation for 723 genes common to both capture kits. Pearson’s correlations were 0.93 (PMT) and 0.92 (SC1) between Kinome and All-Exon captures for RNA from the fresh frozen samples and 0.91 and 0.89 for the FFPE RNA samples, respectively (Fig 2A).

**Fig 2.**
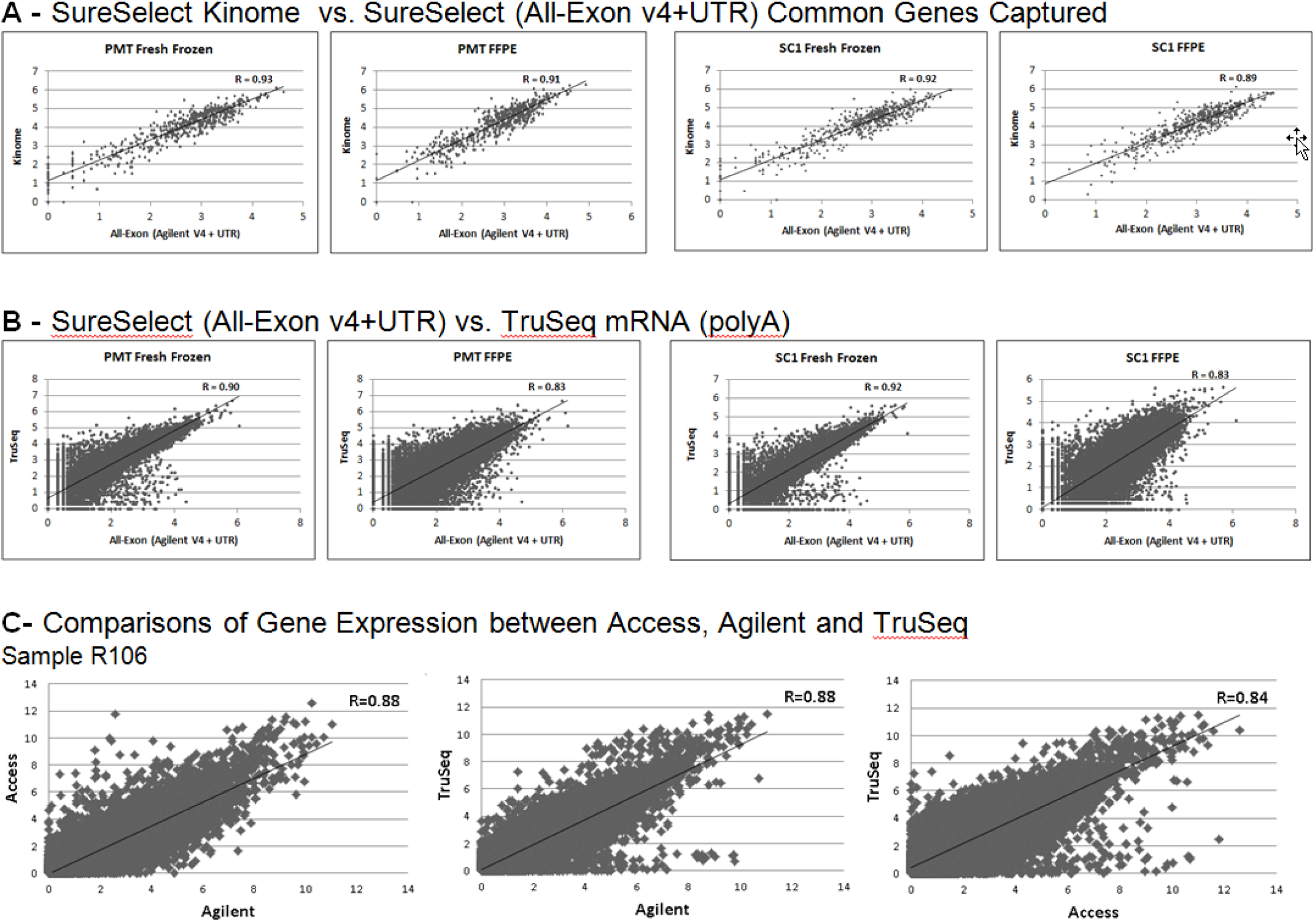
Gene Expression Correlation Comparisons for Different Capture Methods. A) Gene expression correlation plots for shared genes captured by both the Kinome and All-Exon capture methods. B) Gene expression correlation plots for the All-Exon capture and Illumina TruSeq mRNA on RefSeq genes that are included in the All Exon v4+UTR capture bait. C) Gene expression correlation plots for matched FF and FFPE RNA from sample R106 for genes common to TruSeq (FF poly A), Agilent SureSelect All-Exon v6 + UTR bait (FFPE capture) and the Access bait (FFPE capture). Axis scales represent log2 transformed expression value

### Whole Transcriptome Analyses Comparing All-Exon Capture and TruSeq RNA Access vs. mRNA Truseq

When comparing the performance of the All-Exon capture to TruSeq mRNA, a correlation of 0.90 (PMT) and 0.92 (SC1) was observed for gene expression correlation between the two methods for high quality FF RNA samples. Not unexpectedly, gene expression correlation was lower for FFPE RNA with correlations of 0.83 for both the PMT and SC1 samples. (Fig 2B).

To further examine RNA capture with RNA isolated from FFPE tissues, we examined samples containing known fusions using FF RNA analyzed by TruSeq mRNA and the matched FFPE RNA analyzed by either TruSeq RNA Access or SureSelect All-Exon capture. RNA-seq using the Access protocol generally resulted in higher reads mapped to exon-exon junctions (38.2% on average) compared with those by TruSeq mRNA or SureSelect with junction reads averaging 23.6% and 24%, respectively (S Table 6).

We observed significant read differences among the different RNA-seq methods (S Table 6). In addition, we also observed that the method of RNA extraction appeared to affect the average percentage of mapped and junction reads. With the Access protocol, the junction reads averaged 33.6% and 27.1% for the RecoverAll (Ambion) and RNeasy (Qiagen) kits, respectively, compared to 17% and 15% for the SureSelect capture kit (S Table 6). For sample R106, gene expression correlation (R) between TruSeq mRNA using FF RNA and TruSeq RNA Access using the matched FFPE RNA was 0.84. Correlation between TruSeq mRNA (FF) and Agilent SureSelect (FFPE) was slightly higher at 0.88 possibly due to the fact that Agilent SureSelect similarly captures the 3’ untranslated regions and thus more closely resemble poly-A selected mRNA analyzed by the TruSeq mRNA protocol. Correlation between the two capture methods (Illumina Access and Agilent SureSelect for FFPE RNA was the same also at R = 0.88. (Fig 2C).

### Detection of Clinically Significant Fusions Using Capture Based RNAseq Methods

Among the samples tested, fusion detection across the three methods showed an increase in fusion supporting reads using the capture methods compared to TruSeq mRNA with poly-A selection. Among the samples tested, we observed no fusion supporting reads in the non tumor sample (R106) by all three methods (Table 4). Fusions were detected by TruSeq mRNA sequencing for FF sample R152 but not R130. In contrast, both capture-based methods were able to identify the known fusions in these two FF samples, with Access having 11-605 supporting reads compared to SureSelect with 4-121 supporting reads (Table 4A). The failure of TruSeq mRNA to detect a fusion in sample R130 is most likely due to the distance of the *LPP-FOXP1*(>5kb) and *FOXP1-LPP* (>15kb) fusions from the 3’ poly-A site as we have previously shown that poly-A selection based mRNA-seq is less likely to detect gene fusions that are distant from the 3’ end of the mRNA (3).

Same as the results from FF RNA, no fusion reads were found in the matching R106 FFPE sample. The same *LPP-FOXP1* and *FOXP1-LPP* fusions were identified by the Access protocol even though the *FOXP1-LPP* fusion was >15kb away from the 3’ end of the gene. For sample R153, both fusions were identified by the capture methods, with the Access protocol consistently having more supporting reads than the SureSelect protocol (Table 4B). Although not a quantitative test, read depth also likely reflects the relative level of fusion gene expression in the sample (S Table 7).

## Discussion

When comparing FFPE RNA isolation methods, we found that the Ambion RecoverAll method showed consistent results across isolations as well as allowed for recovery of both RNA and DNA. The Qiagen miRNeasy method gave more variable DV_200_ values and lower RNA yields, but had a higher purity of RNA which is also an important factor to consider for RNA-seq analysis. Despite these differences, it was assuring to observe that when RNA samples had DV_200_ values greater or equal to 30%, gene expression measurements by RNA-seq were robust and comparable using either TruSeq Stranded Total RNA or the capture based methods (Figure 1, Figure 2A). As the DV_200_ value decreased (Qiagen miRNeasy FFPE Liver Tumor), increasing the input amount of RNA was found to improve the robustness of the assay and generate comparable read statistics (S Table 3).

Similar to other reports (8, 9, 14) we observed that RNA capture methods were highly effective at pulling down the targeted transcripts. This strategy enabled greater than 70% of reads to map to the targeted genes and exon-exon junctions thus enabling a strong gene expression correlation and fusion gene detections in FFPE RNA at levels comparable to or better than with poly-A selection based TruSeq mRNA sequencing despite the fact that FFPE RNA samples are high degraded and FFPE RNA libraries often have shorter insert size relative to the matched fresh frozen libraries.

Although mRNA TruSeq using RNA from high quality fresh frozen tissues gave better correlation than capture-based method for gene expression analysis (Fig.2B), the sensitivity of fusion detection markedly decreased for lower quality RNA samples for gene fusion junctions further away from the 3’ end of the gene (3). Therefore, it is worth considering gene capture based methods even for RNA from fresh frozen tissues (4), especially for cases with low RIN values, since these samples had the highest percentage of reads mapped to genes when subjected to capture with the All-Exon bait. Furthermore, our results showed that whole transcriptome based capture on FFPE samples could generate sequencing data that correlates well to TruSeq mRNA despite the significant differences in tissue fixation, sample preparation, RNA quality, and library construction methods.

Our study also showed that selective capture of smaller regions such as the Kinome yielded a higher percentage of exonic base coverage versus a whole transcriptome capture (S Figure 1). This increased depth of coverage is likely to increase the sensitivity of fusion detection and improve the discovery of novel fusions in the targeted genomic areas of interest as well as the presence of transcript isoforms and expressed somatic variants. The high correlation between Kinome and the All-Exon Capture demonstrates that gene expression levels are still reproducibly measured with both methods despite the saturation of sequencing reads for the less complex target regions. Similar results were observed for RNA from FF as well as FFPE tissues further indicating that the capture based approach for RNA from FFPE tissue is an effective, reliable method for targeted or overall gene expression analysis and fusion detection by RNA-seq.

Lastly, we observed differences in capture efficiency between the Agilent SureSelect All-Exon v6 + UTR and the Illumina Access capture kits. This is at least in part due to differences in the bait design of the different kits. The Agilent SureSelect All-Exon v6 +UTR baits frequently have probes located in the intronic region of the genes in addition to the 3’ UTR due to their primary purpose to capture DNA for sequencing. In contrast, the Access baits do not extend to the 3’ UTR and are primarily located at the coding transcript regions (S Figure 2). A more focused design of Agilent baits to only genes of clinical interest will likely reduce these performance differences to increase its sensitivity for fusion detection.

In conclusion, we show through comprehensive evaluation that there are significant differences in RNA yield and purity among commonly used FFPE RNA extraction methods. However, when RNA samples of sufficient quality and input are used, comparable whole transcriptome NGS profiles can be obtained. Furthermore, gene capture based approach is a highly reproducible and efficient method for whole transcriptome, targeted gene expression profiling as well as known and novel fusion detection using archival FFPE or partially degraded RNA from diverse clincial samples.

## Supporting information

Supplemental Files_Hilker-etal

## Acknowledgements

This work was supported in part by the Mayo Clinic Center for Individualized Medicine and the Comprehensive Cancer Center to the Genome Analysis Core, as well as the Collaborative Research Funds to JJ and KCH from the Department of Laboratory Medicine and Pathology. We are in debt to members of the Genome Analysis Core and to Drs. Wieben and Cunningham for their support through the course of this study and to members of the Mayo Clinic Biospecimen Accessioning and Processing Core for supporting the RNA extraction method evaluations involved in this study. Current address for JJ is Celgene Corporation, 10300 Campus Point Drive, San Diego, CA 92121

## Compliance with ethical standards

### Conflict of interest

The authors declare no conflict of interest.

### Supplementary information is available

## References

1. Lodish H, Arnold B, Zipursky SL, et al. Molecular Cell Biology, 4th edition. New York: W. H Freeman; 2000.

2. Adiconis X, Borges-Rivera D, Satija R, et al. Comparative analysis of RNA sequencing methods for degraded or low-input samples. Nat Methods. 2013;10(7):623–9.

3. Davila JI, Fadra NM, Wang X, et al. Impact of RNA degradation on fusion detection by RNA-seq. BMC Genomics. 2016;17(1):814.

4. Winter JL, Davila JI, McDonald AM, et al. Development and Verification of an RNA Sequencing (RNA-Seq) Assay for the Detection of Gene Fusions in Tumors. J Mol Diagn. 2018;Jul20(4):495–511.

5. Penland SK, Keku TO, Torrice C, et al. RNA expression analysis of formalin-fixed paraffin-embedded tumors. Lab Invest. 2007;87(4):383–91.

6. Cui P, Lin Q, Ding F, et al. A comparison between ribo-minus RNA-sequencing and polyA-selected RNA-sequencing. Genomics. 2010;96(5):259–65.

7. Esteve-Codina AE, Arpi O, Martinez-Garcia M, et al. A Comparison of RNA-Seq Result from Paired Formalin-Fixed Paraffin-Embedded and Fresh-Frozen Glioblastoma Tissue Samples. PLoS One. 2017: 18.

8. Mercer TR, Clark MB, Crawford J, et al. Targeted sequencing for gene discovery and quantification using RNA CaptureSeq. Nat Protoc. 2014;9(5):989–1009.

9. Cieslik M, Chugh R, Wu Y, et al. The use of exome capture RNA-seq for highly degraded RNA with application to clinical cancer sequencing. Genome Res. 2015;25(9):1372–81.

10. Kalari KR, Nair AA, Bhavsar JD, et al. MAP-RSeq: Mayo Analysis Pipeline for RNA sequencing. BMC Bioinformatics 2014, 15:224.

11. Trapnell C, Pachter L, Salzberg SL. TopHat: discovering splice junctions with RNA-Seq. Bioinformatics 2009, 25:1105–1111.

12. Kim D, Salzberg SL. TopHat-Fusion: an algorithm for discovery of novel fusion transcripts. Genome Biol. 2011, 12:R72.

13. Thorvaldsdottir H, Robinson JT, Mesirov JP. Integrative Genomics Viewer (IGV): high-performance genomics data visualization and exploration. Brief Bioinform 2013, 14:178–192.

14. Mercer TR, Gerhardt DJ, Dinger ME, et al. Targeted RNA sequencing reveals the deep complexity of the human transcriptome. Nat Biotech. 2012;30(8):99–106.

