## Supplemental Files_Hilker-etal for "Impact of RNA Extraction and Target Capture Methods on RNA Sequencing Using Formalin-Fixed, Paraffin Embedded Tissues"

### Supplementary Materials

#### Supplemental Methods

Library preparations were done using the following kits:

TruSeq RNA Sample Preparation v2 protocol (Part # 15026495 Rev. F),

TruSeq Stranded mRNA Sample Preparation Guide (Part # 15031047 Rev. E),

TruSeq Stranded Total RNA Sample Preparation Guide (Part # 15031048 Rev. E),

NEBNext Ultra RNA Library Preparation (NEB Instruction manual #E7530S/L),

TruSeq RNA Access Library Prep Guide (Part # 15049525 Rev. B),

SureSelect$^{XT}$ RNA Target Enrichment for Illumina Multiplexed Sequencing Strand-Specific RNA Library Prep and Target Enrichment Protocol (Version D0) was used with modifications for FFPE Derived Total RNA (Application Note - Siebold, Arezi, 2016).
Available from: http://cn.agilent.com/cs/library/datasheets/public/FFPE%20RNA-Seq%20App%20Note%205991-4794EN.pdf.

#### Supplementary Data Sheet Publications


Sooknanan R, Pease J, Doyle K. Novel methods for rRNA removal and directional, ligation-free RNA-seq library preparation. Advertising Feature-Application note. Nat Methods. © Nature America Inc. 2010.

TruSeq RNA Access Library Prep Kit. Data sheet: RNA Sequencing. © Illumina Inc. 2014.

Siebold AA, Bahram A. Utilization of FFPE Derived Total RNA for Targeted or Whole Transcriptome RNA-seq using SureSelect Strand, Application note. © Agilent Technologies, Inc. 2014, 2016.

TruSeq RNA Exome. data sheet © Illumina Inc. 2017.

Jones CJ, Siebold AA, Lucus AB. Enrichment and Ribosomal Depletion of FFPE RNA. Application note. © Agilent Technologies, Inc. 2017.

### Supplemental Tables and Figures

#### S Table 1 – Summary of Capture Baits

| Company | Capture Bait Name | Probes | RefSeq Exome Representation (%) | Number of Genes Targeted | Number of Exons Targeted | Size of Targeted Region |
|---|---|---|---|---|---|---|
| Illumina | TruSeq Access | 425,000 | 98 | 21,415 | 214, 126 | 45.0 Mb |
| Agilent | SureSelect All Exon v4 + UTR | 789,000 | 99 | 20,965 | 335,765 | 71.5 Mb |
| Agilent | SureSelect All-Exon v6 + UTR | 1,120,387 | 99 | 23,519 | 450,580 | 91.0 Mb |
| Agilent | SureSelect Kinome | 102,184 | - | 756 | 10,282 | 3.2 Mb |

#### S Table 2 – FFPE RNA Kit Comparison

Roche High Pure FFPE RNA Micro Kit

| Tissue | Paraffin Block | Elution volume (µL) | ng/µL | Total Yield (µg) | 260/280 | 260/230 | RIN | $DV_{200}$ (%) |
|---|---|---|---|---|---|---|---|---|
| Liver Tumor | RR12-2321 A1 | 20 | 16.0 | 0.3 | 1.59 | 0.63 | 2.5 | 74 |
| Colon Tumor 1 | RR12-1133 A4 | 20 | 206.0 | 4.1 | 1.93 | 1.78 | 2.4 | 72 |
| Colon Tumor 2 | RR12-1133 A1 | 20 | 88.7 | 1.8 | 1.93 | 1.09 | 2.0 | 75 |
| Pancreas Tumor | RR12-1876 A14 | 20 | 140.4 | 2.8 | 1.92 | 0.75 | 2.3 | 65 |
| Kidney Tumor | RR12-859 A6 | 20 | 80.1 | 1.6 | 1.41 | 0.15 | 2.0 | 74 |
| Tonsil Normal | RR14-405 A2 | 20 | 228.0 | 4.6 | 1.94 | 1.91 | 2.4 | 78 |

Ambion RecoverAll Total Nucleic Acid Isolation Kit

| Tissue | Paraffin Block | Elution volume (µL) | ng/µL | Total Yield (µg) | 260/280 | 260/230 | RIN | $DV_{200}$ (%) |
|---|---|---|---|---|---|---|---|---|
| Liver Tumor | RR12-2321 A1 | 60 | 231.3 | 13.9 | 1.97 | 1.71 | 2.5 | 52 |
| Colon Tumor 1 | RR12-1133 A4 | 60 | 217.5 | 13.1 | 2.01 | 2.02 | 2.5 | 78 |
| Colon Tumor 2 | RR12-1133 A1 | 60 | 126.6 | 7.6 | 2.03 | 1.86 | 2.4 | 71 |
| Pancreas Tumor | RR12-1876 A14 | 60 | 170.6 | 10.2 | 1.89 | 1.36 | 2.4 | 45 |
| Kidney Tumor | RR12-859 A6 | 60 | 54.0 | 3.2 | 2.06 | 0.93 | 2.3 | 55 |
| Tonsil Normal | RR14-405 A2 | 60 | 214.0 | 12.8 | 1.94 | 1.79 | 2.4 | 69 |

Qiagen miRNeasy FFPE Kit

| Tissue | Paraffin Block | Elution volume (µL) | ng/µL | Total Yield (µg) | 260/280 | 260/230 | RIN | $DV_{200}$ (%) |
|---|---|---|---|---|---|---|---|---|
| Liver Tumor | RR12-2321 A1 | 20 | 328.5 | 6.6 | 1.88 | 1.74 | 2.6 | 3 |
| Colon Tumor 1 | RR12-1133 A4 | 20 | 620.2 | 12.4 | 1.95 | 1.91 | 2.4 | 66 |

| | | | | | | | | |
|---|---|---|---|---|---|---|---|---|
| Colon Tumor 2 | RR12-1133 A1 | 20 | 150.3 | 3.0 | 1.95 | 1.94 | 2.4 | 33 |
| Pancreas Tumor | RR12-1876 A14 | 20 | 392.3 | 7.9 | 1.94 | 2.03 | 2.6 | 22 |
| Kidney Tumor | RR12-859 A6 | 20 | 926.9 | 18.5 | 1.93 | 2.07 | 2.6 | 28 |
| Tonsil Normal | RR14-405 A2 | 20 | 261.6 | 5.2 | 1.95 | 1.87 | 2.4 | 33 |

Qiagen miRNeasy FFPE Kit automated on Qiacube

| Tissue | Paraffin Block | Elution volume (µL) | ng/µL | Total Yield (µg) | 260/280 | 260/230 | RIN | $DV_{200}$ (%) |
|---|---|---|---|---|---|---|---|---|
| Liver Tumor | RR12-2321 A1 | 20 | 550.7 | 11.0 | 1.98 | 1.89 | 2.5 | 65 |
| Colon Tumor 1 | RR12-1133 A4 | 20 | 189.2 | 3.8 | 1.9 | 1.79 | 2.6 | 46 |
| Colon Tumor 2 | RR12-1133 A1 | 20 | 220.4 | 4.4 | 1.96 | 1.87 | 2.5 | 52 |
| Pancreas Tumor | RR12-1876 A14 | 20 | 364.3 | 7.3 | 1.96 | 1.94 | 2.4 | 54 |
| Kidney Tumor | RR12-859 A6 | 20 | 161.4 | 3.2 | 1.89 | 1.61 | 2.3 | 44 |
| Tonsil Normal | RR14-405 A2 | 20 | 297.6 | 6.0 | 1.94 | 2.03 | 2.4 | 41 |

Qiagen RNeasy FFPE Kit

| Tissue | Paraffin Block | Elution volume (µL) | ng/µL | Total Yield (µg) | 260/280 | 260/230 | RIN | $DV_{200}$ (%) |
|---|---|---|---|---|---|---|---|---|
| Liver Tumor | RR12-2321 A1 | 20 | 170.2 | 3.4 | 1.92 | 1.69 | 2.4 | 14 |
| Colon Tumor 1 | RR12-1133 A4 | 20 | 325.5 | 6.5 | 1.98 | 2.10 | 2.6 | 59 |
| Colon Tumor 2 | RR12-1133 A1 | 20 | 385.1 | 7.7 | 1.93 | 1.88 | 2.3 | 50 |
| Pancreas Tumor | RR12-1876 A14 | 20 | 387.2 | 7.7 | 1.97 | 2.07 | 2.6 | 38 |
| Kidney Tumor | RR12-859 A6 | 20 | 410.5 | 8.2 | 1.96 | 1.99 | 2.4 | 29 |
| Tonsil Normal | RR14-405 A2 | 20 | 532.5 | 10.7 | 1.95 | 2.04 | 2.4 | 37 |

**S. Table 3 – RNA-seq Read Statistics for FFPE RNA Extraction Comparison**

| Sample | Total Reads | Used Reads | Mapped Reads (%) | Genome Reads (%) | Junction Reads (%) | Gene Count (%) |
|---|---|---|---|---|---|---|
| High Pure Liver Tumor | 45,895,486 | 45,509,695 | 38,243,266 (83.3) | 34,434,729 (75.0) | 3,808,537 (8.3) | 10,196,247 (22.2) |
| High Pure Colon Tumor 1 | 59,933,354 | 59,875,409 | 51,246,777 (85.5) | 48,241,445 (80.5) | 3,005,332 (5.0) | 15,198,636 (25.4) |
| High Pure Colon Tumor 2 | 73,852,852 | 73,791,250 | 63,351,116 (85.8) | 57,537,645 (77.9) | 5,813,471 (7.9) | 20,702,814 (28.0) |
| High Pure Pancreas Tumor | 61,293,154 | 61,243,597 | 52,005,134 (84.8) | 49,449,129 (80.7) | 2,556,005 (4.2) | 12,814,138 (20.9) |
| High Pure Kidney Tumor | 72,019,140 | 71,954,102 | 61,799,883 (85.8) | 57,023,803 (79.2) | 4,776,080 (6.6) | 19,519,373 (27.1) |
| High Pure Tonsil Normal | 58,433,532 | 58,359,807 | 45,595,145 (78.0) | 42,692,053 (73.1) | 2,903,092 (5.0) | 12,670,145 (21.7) |
| miRNeasy Liver Tumor | 41,894,270 | 41,614,148 | 31,099,257 (74.2) | 29,255,942 (69.8) | 1,843,315 (4.4) | 4,982,537 (11.9) |
| miRNeasy Colon Tumor 1 | 77,659,822 | 77,584,367 | 66,304,021 (85.4) | 62,434,115 (80.4) | 3,869,906 (5.0) | 19,024,949 (24.5) |
| miRNeasy Colon Tumor 2 | 68,920,472 | 68,835,684 | 55,389,226 (80.4) | 50,451,031 (73.2) | 4,938,195 (7.2) | 16,325,977 (23.7) |
| miRNeasy Pancreas Tumor | 73,345,024 | 73,266,018 | 57,018,980 (77.7) | 54,098,837 (73.8) | 2,920,143 (4.0) | 13,514,666 (18.4) |
| miRNeasy Kidney Tumor | 97,491,748 | 97,394,984 | 81,110,043 (83.2) | 73,984,879 (75.9) | 7,125,164 (7.3) | 25,506,294 (26.2) |
| miRNeasy Tonsil Normal | 55,242,686 | 55,097,642 | 38,249,264 (69.2) | 35,193,395 (63.7) | 3,055,869 (5.5) | 10,637,495 (19.3) |
| RecoverAll Liver Tumor | 53,686,378 | 53,665,613 | 43,500,152 (81.0) | 38,778,852 (72.2) | 4,721,300 (8.8) | 13,131,532 (24.5) |
| RecoverAll Colon Tumor 1 | 71,979,760 | 71,967,159 | 59,767,756 (83.0) | 55,669,025 (77.3) | 4,098,731 (5.7) | 18,079,139 (25.1) |
| RecoverAll Colon Tumor 2 | 60,933,016 | 60,919,855 | 51,338,823 (84.3) | 46,590,494 (76.5) | 4,748,329 (7.8) | 16,592,662 (27.2) |
| RecoverAll Pancreas Tumor | 65,014,090 | 65,004,827 | 55,034,541 (84.7) | 51,983,816 (80.0) | 3,050,725 (4.7) | 13,955,566 (21.5) |
| RecoverAll Kidney Tumor | 78,708,774 | 78,698,930 | 67,961,536 (86.3) | 60,422,519 (76.8) | 7,539,017 (9.6) | 27,286,195 (34.7) |
| RecoverAll Tonsil Normal | 73,856,738 | 73,824,853 | 61,614,980 (83.4) | 57,893,270 (78.4) | 3,721,710 (5.0) | 15,257,814 (20.7) |

**S Table 4 – Gene Coverage for T47D+ and T47D- samples with and without capture**

| Samples | T47D+ | | T47D- | |
|---|---|---|---|---|
| | Agilent SureSelect All-Exon v4 + UTR Capture | No Capture | Agilent SureSelect All-Exon v4 + UTR Capture | No Capture |
| **# Genes with ≥ 1X coverage** | 17,386 | 17,625 | 17,686 | 17,958 |
| **# Genes with ≥ 5X coverage** | 15,981 | 16,066 | 16,340 | 16,361 |

Gene  coverage of the two samples with and without capture (TruSeq stranded mRNA) at non-zero coverage and greater than 5X coverage.

**S Table 5 – Read Statistics for matching fresh frozen and FFPE RNA samples of PMT and SC1 for TruSeq mRNA and SureSelect RNA capture**

| | Sample Type | Total Reads | % Mapped | % Mapped to Genome | % Mapped to Junctions | % Mapped To Genes |
|---|---|---|---|---|---|---|
| | Fresh Frozen TruSeq mRNA | 36,411,832 | 96.4 | 80.8 | 15.6 | 81.7 |
| | Fresh Frozen All-Exon Capture | 35,757,261 | 96.2 | 64.6 | 31.6 | 82.1 |
| **PMT** | Fresh Frozen Kinome Capture | 35,685,110 | 95.9 | 65.9 | 30 | 73.2 |
| | FFPE All Exon Capture | 36,155,495 | 92.9 | 67.8 | 25.1 | 78.2 |
| | FFPE Kinome Capture | 36,207,173 | 92.9 | 66.5 | 26.4 | 77.6 |
| | Fresh Frozen TruSeq mRNA | 36,320,412 | 96.4 | 78.7 | 17.9 | 93.3 |
| | Fresh Frozen All-Exon Capture | 35,690,330 | 96 | 63.3 | 32.7 | 79.1 |
| **SC1** | Fresh Frozen Kinome Capture | 35,931,232 | 95.9 | 63.8 | 32.1 | 77 |
| | FFPE All-Exon Capture | 36,026,128 | 93.5 | 65.1 | 28.4 | 76.6 |
| | FFPE Kinome Capture | 36,087,109 | 93.8 | 65.9 | 28 | 80.9 |

Read statistics for samples PMT and SC1 comparing total RNA capture with All-Exon and Kinome baits on matched FFPE and fresh frozen derived RNA. As a standard, the read statistics from Illumina TruSeq on fresh frozen derived RNA was included. In each case, the percentage was calculated based on the genes each kit targets. (Kinome = 756 genes, All -Exon = 20,002 genes and for TruSeq the Ref Seq transcriptome).
% mapped represents the total percentage of reads mapped to the genome + transcriptome.
% mapped to Genome is the percentage of reads mapped to contiguous regions in the genome which don't have any exon-exon or exon-intron junctions.
% mapped to Junctions is the percentage of reads that map to exon-exon/exon-intron junctions i.e they map to transcriptome.
% Mapped to Genes shows the total percent of reads mapping within gene bodies.

**S Fig 1 – Violin Plot Comparisons of Exonic Base Coverage for Al- Exon and Kinome Capture for FFPE samples PMT and SC1**

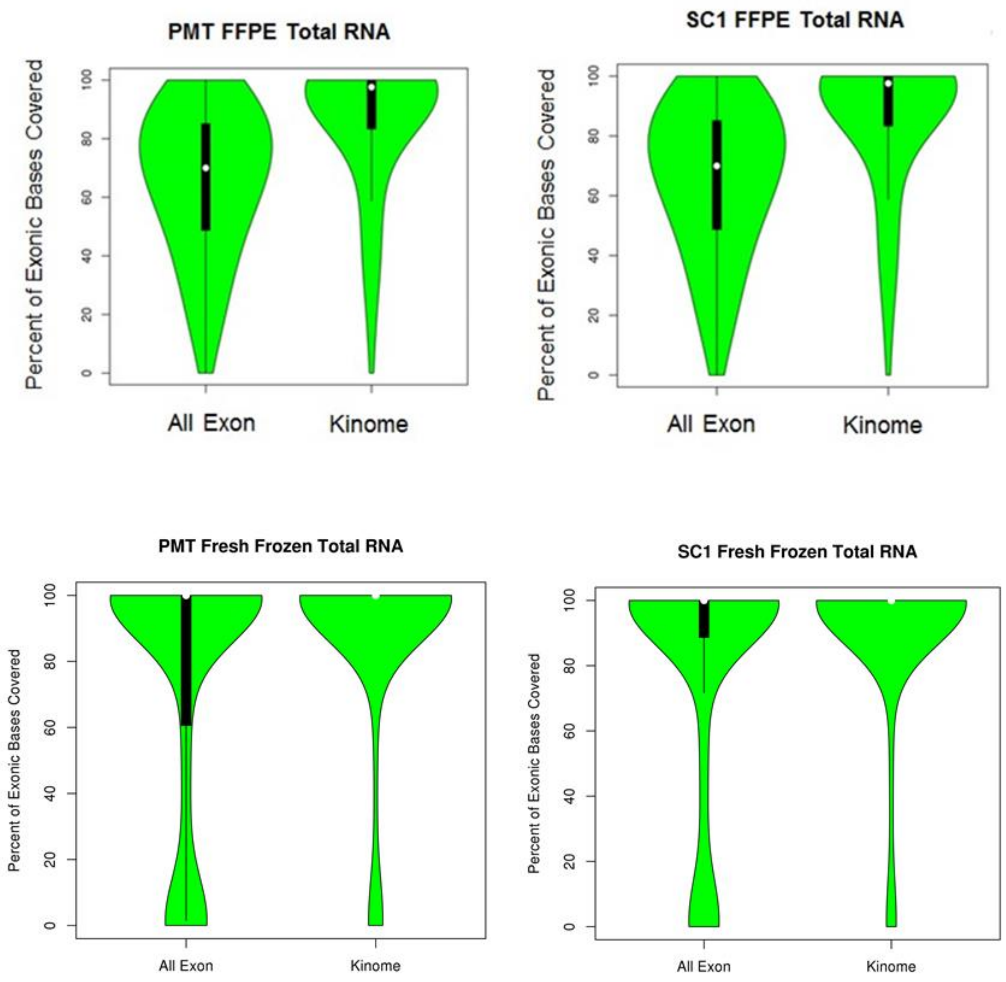


Violin plots representing the exonic base coverage for all-exon capture and kinome capture on commonly captured regions for samples PMT and SC1. The kinome capture shows a higher percentage of reads being made up of exonic bases versus All-Exon capture.

**S Table 6 – RNA-seq Read Comparison for Fresh Frozen and FFPE RNA Samples Captured with TruSeq*, Access** or SureSelect*****

| Tissue and Capture Type | Number of Samples | Average Mapped Reads (%) | Average Mapped Junction Reads (%) |
|---|---|---|---|
| **Fresh Frozen** | | | |
| TruSeq | 3 | 93.2 | 23.6 |
| Access | 3 | 88.2 | 38.2 |
| SureSelect | 3 | 87.0 | 24.0 |
| **FFPE (Ambion RecoverAll)** | | | |
| Access | 3 | 80.9 | 33.6 |
| SureSelect | 3 | 73.0 | 17.0 |
| **FFPE (Qiagen miRNeasy)** | | | |
| Access | 3 | 66.8 | 27.1 |
| SureSelect | 3 | 69.0 | 15.0 |

*TruSeq – PolyA selection done before library preparation.
**Access – Illumina Access baits used for capture during library preparation.
***SureSelect – Agilent SureSelect All-Exon v6 + UTR baits used for capture during library preparation.

**S Table 7 Mapped Reads used in Fusion Detection in Fresh Frozen and FFPE Tissue using TruSeq mRNA, Access Capture and SureSelect Capture**

| Tissue Type | Assay | Sample | Total Reads | Used Reads | Mapped Reads (%) | Mapped Genome Reads (%) | Mapped Junction Reads (%) |
|---|---|---|---|---|---|---|---|
| Fresh Frozen | TruSeq | R106 | 89,550,435 | 89,537,716 | 83,315,110 (93.0) | 65,605,295 (73.3) | 17,709,815 (19.8) |
| | | R130 | 95,616,276 | 95,602,972 | 91,300,001 (95.5) | 60,613,355 (63.4) | 30,686,646 (32.1) |
| | | R152 | 117,471,572 | 117,438,144 | 111,312,221 (94.8) | 94,478,375 (80.4) | 16,833,846 (14.3) |
| | Access | R106 | 58,595,412 | 58,585,552 | 53,846,928 (91.9) | 30,772,505 (52.5) | 23,074,423 (39.4) |
| | | R130 | 64,234,444 | 64,224,170 | 58,797,098 (91.5) | 35,218,831 (54.8) | 23,578,267 (36.7) |
| | | R152 | 70,801,972 | 70,791,415 | 65,067,629 (91.9) | 35,677,715 (50.4) | 29,389,914 (41.5) |
| | SureSelect | R106 | 47,607,506 | 47,400,990 | 40,897,969 (85.9) | 30,541,037 (64.2) | 10,356,932 (21.8) |
| | | R130 | 53,803,494 | 53,565,427 | 47,125,617 (87.6) | 33,836,291 (62.9) | 13,289,326 (24.7) |
| | | R152 | 54,401,686 | 54,164,953 | 48,020,187 (88.3) | 34,389,923 (63.2) | 13,630,264 (25.1) |
| FFPE | Access | R106 | 38,538,134 | 38,526,864 | 32,358,744 (84.0) | 18,158,823 (47.1) | 14,199,921 (36.8) |
| | | R130 | 54,769,710 | 54,753,933 | 47,335,133 (86.4) | 29,245,942 (53.4) | 18,089,191 (33.0) |
| | | R153 | 53,341,192 | 53,325,712 | 45,633,673 (85.6) | 27,738,451 (52.0) | 17,895,222 (33.5) |
| | SureSelect | R106 | 61,641,810 | 61,632,684 | 43,998,952 (71.4) | 33,996,052 (55.2) | 10,002,900 (16.2) |
| | | R130 | 68,539,714 | 68,528,435 | 51,575,116 (75.2) | 39,317,678 (57.4) | 12,257,438 (17.9) |
| | | R153 | 65,807,100 | 65,797,907 | 48,107,331 (73.1) | 37,799,209 (57.4) | 10,308,122 (15.7) |

#### S Figure 2 – Probe Design Impact on Capture

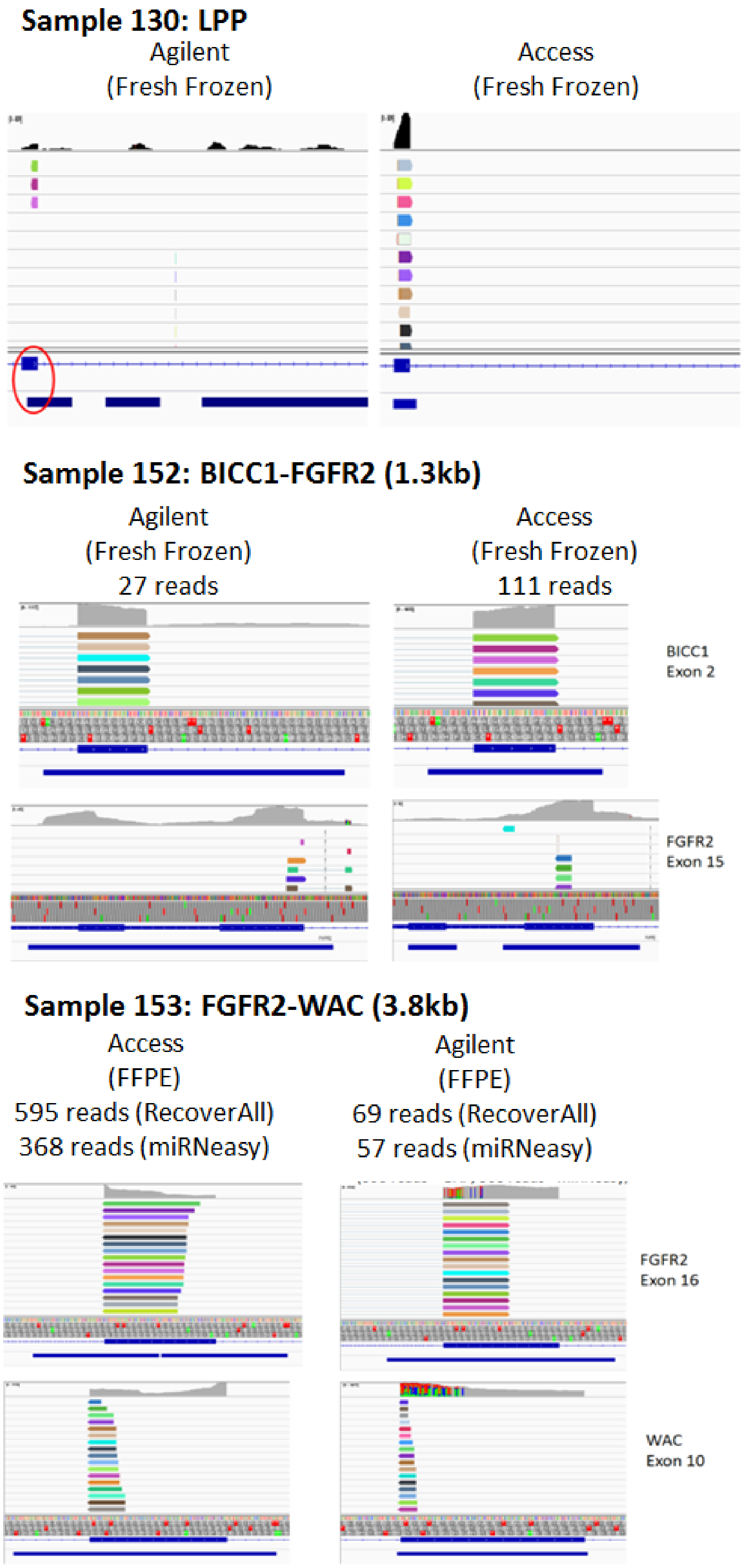


Examples of read coverage differences at gene/exon locations, as indicated, by Access (left panels) or Agilent (right panel) baits. Blue bars under each IGV plot indicate regions of bait location.
